# Bayesian Multivariate Growth Mixture Modeling of Longitudinal Data: An Application to Alzheimer’s Disease Study

**DOI:** 10.1101/2021.03.10.434854

**Authors:** Wenyi Lin, Michael C. Donohue, Philip Insel, Armin Schwartzman, Wesley K. Thompson

**Affiliations:** Division of Biostatistics, Department of Family Medicine and Public Health, University of California San Diego, La Jolla, CA, USA; Alzheimer’s Therapeutic Research Institute,University of Southern California, San Diego,CA, USA; Clinical Memory Research Unit, Department of Clinical Sciences Malmö, Lund University, Sweden; Halıcıoğlu Data Science Institute, University of California San Diego, La Jolla, CA, USA

## Abstract

Alzheimer’s disease (AD) studies often collect longitudinal biomarker measures of multiple cohorts at different stages of disease and follow these biomarkers with a relatively short period of time. The heterogeneity of the longitudinal patterns of biomarkers can be ubiquitous across both individual trajectories and cognitive domains. We propose a flexible Bayesian multivariate growth mixture model to identify distinct longitudinal patterns of data from the Alzheimer’s Disease Neuroimaging Initiative (ADNI) study. A Gibbs sampling is implemented for achieving the Bayesian inference. We perform a simulation study to demonstrate the adequate performance of our proposed approach and apply the model to identify three latent cognitive decline patterns among patients from the ADNI study.

## 1 Introduction

Alzheimer’s disease (AD), an irreversible neurodegenerative disorder, is the most common cause of dementia, affecting millions across the world. Biomarkers such as those derived from various neuroimaging modalities are commonly used to characterize progression of AD pathology. Tracking the temporal evolution of specific AD biomarkers may improve understanding of disease mechanisms and thus indicateds the onset and progression of clinical symptoms[1, 2]. Recent studies have emphasized the importance of using biomarkers to predict risk of decline from mild cognitive impairment to dementia[3, 4] and the modeling of AD biomarker trajectories can be used to predict future clinical course, including cognitive and functional decline. Therefore, prediction of AD progression is potentially valuable in clinical practice, as preventive measures can be more effective prior to onset of severe AD symptoms, i.e. when patients are cognitively normal (CN) or mildly cognitively impaired (MCI) [5, 6].

Mixed-effects models provide a flexible and powerful statistical framework for analyzing longitudinal biomarker data that enables characterization of between-subject variation in within-subject AD progression, accounting for the considerable heterogeneity in individual trajectories. It has been widely implemented with previous AD studies, including applications of magnetic resonance imaging (MRI) measures [7], positron emission tomography (PET) measures [8], and clinical and neuropsychological assessments[9]. However, such heterogeneity of cognitive decline may also vary across multiple sources of biomarkers[1, 2]. Trajectories of multiple biomarkers in union may be sensitive to changing underlying disease states well before cognitive or functional impairments are detected by clinical interviews. Additionally, considering most longitudinal studies of AD are of much shorter duration than the length of time involved in progression of the illness, multivariate long-term dynamics modeling of longitudinal trajectories are of great interest and importance[10]. Analytic approaches are thus needed to account for both biomarker dynamics and change in illness states from the short-term multivariate longitudinal data.

Growth mixture models (GMM)[11], which can be seen as an extension of mixed effects models that incorporates latent class analysis, can be usefully applied in this respect. GMMs have been implemented in multiple longitudinal studies related with Alzheimer’s disease. For example, Pietrzak et al.[12] applied GMMs to reveal three predominant trajectories of a composite score of episodic memory change. Lin et al.[13] fit two separate GMMs to examine the potentially heterogeneous longitudinal trajectories of episodic memory and executive function for identifying the existence of successful cognitive agers. In addition, Leoutsakos et al.[14] implemented parallel-process growth mixture models on both cognitive and functional measures and a follow-up multinomial logistic regression to predict class membership in univariate models. These applications either focused on univariate modeling or analyzing post-hoc effects of multiple outcomes based on GMM results. Lai et al.[15] extended previous works and constructed a multivariate finite mixture latent trajectory model, which can identify subgroups of patients. However, the specification of the model still requires constraints with the covariance matrices and the expectation–maximization (EM) algorithm was applied for parameter estimation, whose performance often depends on the starting values.

To address these limitations, we propose a fully Bayesian multivariate growth mixture model (BMGMM) for the analysis of predicting latent states of AD progression, which meanwhile indicates direction of future development of disease based on joint multivariate modeling. Our proposed methodology has several extensions compared with previous work in a number of ways. First, our model enables statistical inferences for modeling multivariate longitudinal growth trajectories and simultaneously predicts latent clusters, incorporating varying random effects across both latent states and outcomes. Second, the proposed model accommodates covariates for both outcome trajectory modeling and latent class modeling. Third, the parameter estimation in a Bayesian framework utilizes several novel applications, considering the complexity of both data and model structure. Specifically, we incorporated a Half-t prior[16] for modeling the class-specific covariance matrix across varying outcomes and the Pólya–Gamma data augmentation approach[17] for facilitating the sampling procedure in modeling the latent state probabilities. Fourth, the proposed model predicted latent states of current disease stage as well as informed a potential future direction of disease progression. Finally, our model is a fully Bayesian hierarchical model which handles longitudinal trajectories of multiple outcomes in a unified modeling framework, allowing flexible and rigorous interrogation of the posterior distribution and inferences about long-term disease dynamics.

## 2 Case Study: Alzheimer’s Disease Neuroimaging Initiative (ADNI) Study

The proposed approach was applied to the Alzheimer’s Disease Neuroimaging Initiative (ADNI) data. ADNI is a multicohort longitudinal study started in 2004, tracking the progression of AD in the human brain with clinical, imaging, genetic and biospecimen biomarkers. Over 1900 volunteers between the ages of 55 and 90 in the stage of cognitively normal or mildly cognitively impaired or AD dementia were recruited and four phases of study has been implemented. The first phase, referred as ADNI-1, consists of 800 individuals: 200 with CN, 400 with MCI, and 200 with mild dementia. Following phases continue enrolling additional individuals at different stages as well as keeping participants from the prior cohorts. More information about the details of the study can be found at http://adni.loni.usc.edu/.

The classification of stages of disease progression in ADNI was given as follows [18]. Individuals in CN have a Clinical Dementia Rating (CDR) score of 0, Mini Mental State Exam (MMSE) between 24-30 and no memory complaints. In addition, the CN subjects could not have any significant impairment in cognitive functions or activities of daily living. The Significant Memory Concern (SMC) cohort was added to the second phase of the study, i.e. ADNI-2. Individuals identified as in the stage of SMC have a self-report significant memory concern, quantified by using the Cognitive Change Index. MCI participants have a CDR of 0.5, MMSE between 24-30, a subjective memory complaint and could not qualify for the diagnosis of dementia. Dementia subjects have CDR of 0.5 or 1, MMSE of 20 to 26, memory complaints and meet criteria for probable AD. At the screening visit, all subjects were required to provide demographics, family history, medical history and physical examinations and neurological examinations were given to record crucial signs.

## 3 Methods

In this section, we developed a Bayesian multivariate growth mixture model (BMGMM) for analyzing multiple outcomes from longitudinal study. Section 3.1 introduces the BMGMM framework combined with a finite mixture model and a multivariate mixed effects model. Section 3.2 and 3.3 provides two novel implementations for valid and efficient Gibbs sampling. Section 3.4, 3.5 and 3.6 illustrate common issues with Bayesian model fitting, accommodated with our model design and data structure.

### 3.1 Model Setting

Let *Y*_*ijl*_ denote the measured outcome *l*, *l* = 1, 2, ..., *L*, for subject *i*, *i* = 1, 2, ..., *n* at time *j*, *j* = 1, ..., *m*_*il*_. Given that subject *i* is in latent class *C*_*i*_ = *k*, *k* = 1, 2, ..., *K*, the mixture latent class mixed model of ***y***_*i*_ has the form

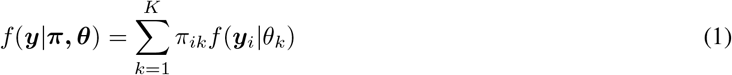

where *θ*_*k*_ is the class-specific parameter for each latent class *k* (*k* = 1, 2*, .., K*). *π*_*ik*_ is the class proportion of each subject *i* and satisfies 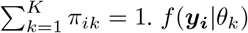 are individual density functions for each latent class and assumed to be normally distributed with the form

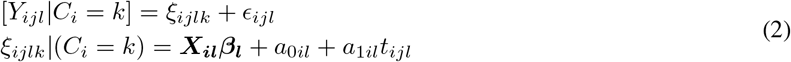

where the fixed effects 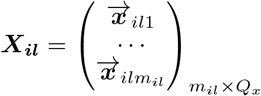 can be either time-varing or time-constant covariates and 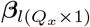 is the corresponding regression coefficient for outcome 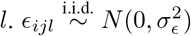 is a vector of error items.

The parameters *a*_0*il*_ and *a*_1*il*_ correspond to subject-specific random intercept and slope for outcome *l*. Let ***a***_***i***_ = (*a*_0*i*1_, *a*_1*i*1_, ..., *a*_0*iL*_, *a*_1*iL*_)′ and ***a***_***i***_ follows a multivariate normal distribution, with the form as ***a***_***i***_|*C*_*i*_ = *k* ~ *N* (***α***_***k***_, Σ_*k*(2*L×*2*L*)_). Here, ***α***_***k***_ are class-specific random effects for all outcomes and Σ_*k*_ are class-specific covariance matrices of random intercepts and slopes among all outcomes.

The class probability ***π*** = (*π*_*i*1_, ..., *π*_*iK*_)′ is modeled using a multinomial logistic regression model:

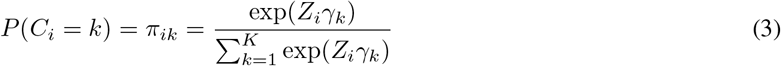

where 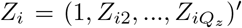 is the class-related covariates and *γ*_*k*_ is the corresponding class-specific coefficient vector. To ensure identifiability of the model, *γ*_*k*_ ≡ 0 when *k* = *K* and it is used as the reference class.

### 3.2 Hierarchical approach for estimating covariance matrix

To accommodate the multivariate extension of modeling the class-specific covariance matrix Σ_*k*_, we adopt Half-t priors [16] to reduce potential impact of misestimation of the correlation coefficients, which is commonly seen when the dimensions of the covariance matrix increase. It can be viewed as an approach of separation strategy and flexibly models variances and correlations components separately.

For class-specific covariance matrix Σ_*k*_, the prior is specified as *Inverse-Wishart*(*ν* + 2*L* − 1, 2*ν*Δ), where Δ is a diagonal matrix with elements *λ*_*l*_, which are assumed to be independently distributed 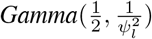. As it is noted in Huang and Wand (2013), we refer this prior as *Half-t*(*ν, **ψ***) for it has been proved that for standard deviations, this prior generates a Half-t distribution with *ν* degrees of freedom and scale parameter *ψ*_*l*_. Specifically, higher value of *ψ*_*l*_ indicates larger uncertainty about the standard deviations. In addition, specification of *ν* = 2 results in uniform prior for the correlation coefficients. The stability of this prior has been proved with our simulation study by comparison with commonly-used Inverse-Wishart priors for covariance matrix.

The full conditional distribution of Σ_*k*_ is given as

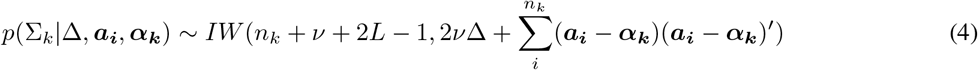

where 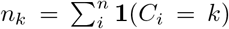 is the number of subjects in latent class *k* and **1**(*C*_*i*_ = *k*) is an indicator function representing subject *i* belongs to class *k*.

### 3.3 Pólya–Gamma data augmentation approach for multinomial regression

For estimating class-specific parameters in multinomial regression, we implement algorithms using a family of Pólya– Gamma distribution, introduced by Polson et al.(2013)[17], and generate the original data-augmentation algorithm for Bayesian logistic regression to multinomial settings. The main strategy of the algorithm is to implement Gaussian draws for generating regression parameters and the Pólya–Gamma draws are incorporated for single layer of latent variables. Compared with previous attempts with missing-data strategy to the logit model[19, 20], which are either approximate or complicated, the Pólya–Gamma method is efficient and simpler.

A random variable *ω* is said to follow Pólya–Gamma distribution with parameters *b >* 0 and *c* ∈ ℝ if

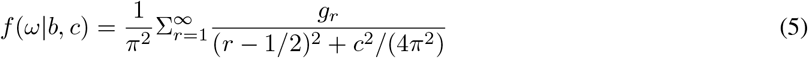

where *, g*_*r*_ ~ *Gamma*(*b,* 1) and *ω* is denoted as *ω* ~ *PG*(*b, c*). Polson et al.(2013)[17] proved that for the Bayesian logistic regression model, the Pólya–Gamma family can yield a simple Gibbs sampler and the posterior distribution is a scale mixture of Gaussians. Based on the derivation procedure for binary outcomes, we extend the algorithm to a multinomial setting.

Suppose that a random variable *ω*_*ik*_ follows Pólya–Gamma distribution with parameters (1, 0), denoted by *PG*(*ω*_*ik*_ |1, 0) and the multinomial model of 3 holds. Following Holmes and Held(2006)[19], the likelihood for *γ*_*k*_ conditional upon *γ*_*−k*_, i.e. the parameter matrix with column vector *γ*_*k*_ removed, is

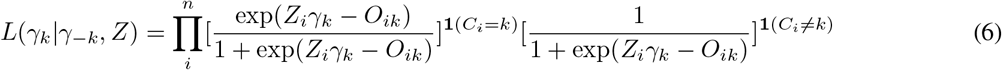

where *O*_*ik*_ = log Σ_*k′*≠*k*_ exp(*Z*_*i*_*γ*_*k*_). Based on the logistic regression form of conditional likelihood, the contribution of *Z*_*i*_ to the likelihood in *γ*_*k*_ can be written as

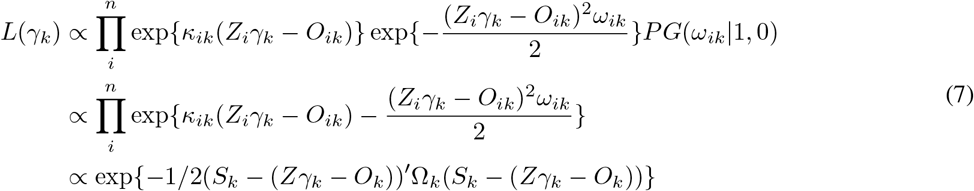

where *κ*_*ik*_ = **1**(*C*_*i*_ = *k*) − 1/2, 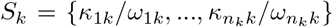, and 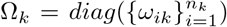. Providing the prior *γ*_*k*_ ~ *N* (*m*_0*k*_, *V*_0*k*_), the posterior is given as

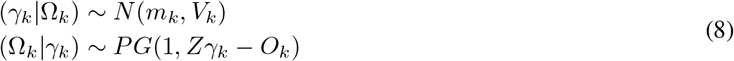

where 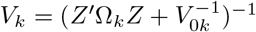 and 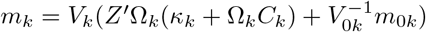. *Z* = {*Z*_1_, ..., *Z*_*n*_} is the aggregated design matrix of class-related covariates and *O*_*k*_ = log Σ_*k′*≠*k*_ exp(*Zγ*_*k*_). Therefore, it allows for Gibbs sampling from the joint posterior distribution without appealing to analytic approximations to the posterior.

### 3.4 Prior specification

The specification of priors is important for Bayesian model fitting since an improper prior can result in an improper posterior. We adopt weakly informative prior distributions to all model parameters. For each ***β***_***l***_ and ***α***_***k***_, an independent conjugate normal distribution 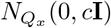 is assigned. Under the Pólya–Gamma sampling scheme, each *γ*_*k*_, *k* = 1, 2, ..., *K* − 1, is assigned a *N* (*m*_0*k*_, *V*_0*k*_) prior. For standard deviations in class-specific covariance matrix Σ_*k*_, Gelman(2006)[21] suggested non-informative Half-t priors that allow for small values. Meanwhile, the prior on the correlation parameter in each Σ_*k*_ is specified as *Uniform*(−1, 1). Finally, for 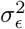, we assume a conjugate *Inverse-Gamma*(*δ*_!_, *δ*_2_) prior.

### 3.5 Posterior Derivation

The introduced prior specification above provides full conditional distributions for all model parameters, which can be efficiently updated via the Gibbs sampler. Assuming prior independence of the model parameters, the joint posterior is given by

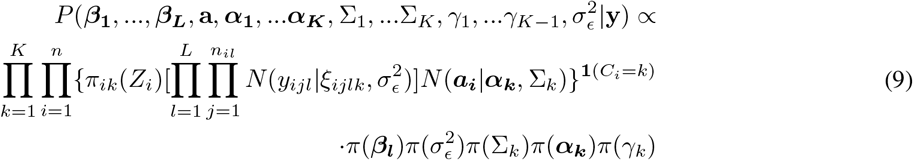

where 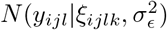 denotes the normal likelihood for *y*_*ijl*_ described in Equation 2. *N* (***a***_***i***_|***α***_***k***_, Σ_*k*_) denotes the normal prior for random effects ***a***_***i***_ with mean ***α***_***k***_ and covariance matrix Σ_*k*_. The rest *π*(·)’s provide the prior distributions for corresponding parameters. Detailed Gibbs sampler can be found in Appendix A.

### 3.6 Model comparison and label switching

We compute average bias, mean squared error and prediction accuracy of latent classes as diagnostic approaches for checking model fitting. We also use the widely applicable information criterion (WAIC) as a way of model comparison criteria[22], which is viewed as an improvement on deviance information criterion (DIC) for Bayesian methods. The WAIC can be computed using the log-likelihood evaluated at the posterior simulations of the parameter values and a lower WAIC is favored in model selection. The R code that provides efficient computation can be found online. The log-likelihood for the whole sample is

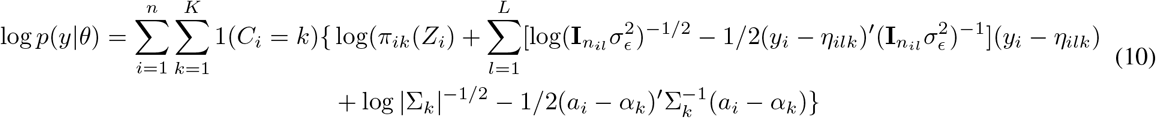

To address the common label switching issue when fitting Bayesian mxiture models, we implemented the Stephens’ method[23] using the `label.switching` package in R[24]. On the other hand, once the truth is known, for both simulation study and ADNI data, the order constraint can be assigned to one parameter in random effects to avoid the potential label switching.

## 4 Simulation study

In simulation study, we conducted experiments for both examining the performance of the proposed algorithm and checking how model misspecification affects the model fitting. For each simulation example, we simulated *n* = 200 individuals and *t* = 6 time points. We kept 75% of the generated data to create a sparser dataset to better mimic the data structure in ADNI study. For each individual, the observation times were sampled from a uniform distribution *t*_*ijl*_ ~ Unif(0, 5) and the baseline time was sampled from a normal distribution *T*_*i*0_ ~ *N* (0, 5) for indicating varying starting time points.

For each model fitting, we ran two parallel Markov chains and each single chain was run 2000 iterations and the first 1000 iterations were discarded as a warm-up phase, yielding a total of 2000 samples for posterior analysis. In addition, each model was simulated *M* = 100 times.

### 4.1 Model fitting with varying numbers of outcomes and latent states

In the first scenario of the simulation study, we simulated examples from *P* = 1, 2, 3 outcomes and *K* = 2, 3 latent states. Let 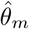 be the posterior estimate of a model parameter *θ*_*m*_ in the *m*-th simulation. The following quantities were considered for assessing the model performance: (i) average bias, 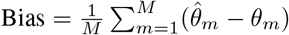, (ii) total mean squared error, 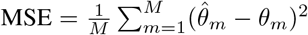 and (ii) the coverage rate of the 95% credible intervals, *C*_95%_. Meanwhile, the prediction accuracy of latent states, *p*_*acc*_, is also included.

Table 1 presents the measures of average bias, MSE and *C*_95_ from one simulation example with *P* = 2 outcomes and *K* = 3 latent classes, including the true simulation parameters. To better mimic the ADNI data, the random intercepts (*a*_0*k*_ = {0.2, 2, 10}) of the first outcome were set to be more separable compared with them of the second outcome (*a*_0*k*_ = {0.5, 1, 2}). Figure 1 plots the coverage rate of the 95% credible intervals of each element in the covariance matrix. Both summary table and plot illustrate that most parameters are estimated well with the proposed model, indicating successful convergence and applicability of the model implementation. Other simulation results from varying combinations of number of latent classes and outcomes are included in Appendix B.

**Table 1.**
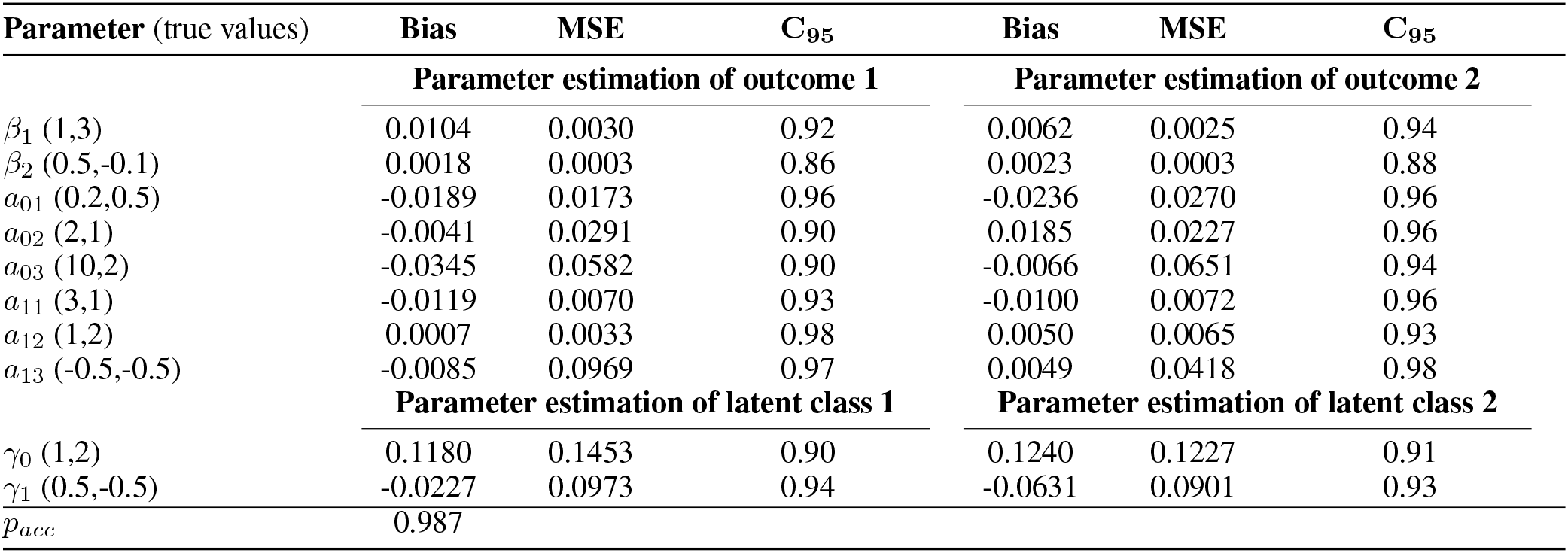
Simulation study results of *P* = 2 and *K* = 3. The model was fit to all 200 samples.

| Parameter (true values) | Bias | MSE | C<sub>95</sub> | Bias | MSE | C<sub>95</sub> |
| --- | --- | --- | --- | --- | --- | --- |
|  | Parameter estimation of outcome 1 |  |  | Parameter estimation of outcome 2 |  |  |
| $\beta_1$ (1,3) | 0.0104 | 0.0030 | 0.92 | 0.0062 | 0.0025 | 0.94 |
| $\beta_2$ (0.5,-0.1) | 0.0018 | 0.0003 | 0.86 | 0.0023 | 0.0003 | 0.88 |
| $a_{01}$ (0.2,0.5) | -0.0189 | 0.0173 | 0.96 | -0.0236 | 0.0270 | 0.96 |
| $a_{02}$ (2,1) | -0.0041 | 0.0291 | 0.90 | 0.0185 | 0.0227 | 0.96 |
| $a_{03}$ (10,2) | -0.0345 | 0.0582 | 0.90 | -0.0066 | 0.0651 | 0.94 |
| $a_{11}$ (3,1) | -0.0119 | 0.0070 | 0.93 | -0.0100 | 0.0072 | 0.96 |
| $a_{12}$ (1,2) | 0.0007 | 0.0033 | 0.98 | 0.0050 | 0.0065 | 0.93 |
| $a_{13}$ (-0.5,-0.5) | -0.0085 | 0.0969 | 0.97 | 0.0049 | 0.0418 | 0.98 |
|  | Parameter estimation of latent class 1 |  |  | Parameter estimation of latent class 2 |  |  |
| $\gamma_0$ (1,2) | 0.1180 | 0.1453 | 0.90 | 0.1240 | 0.1227 | 0.91 |
| $\gamma_1$ (0.5,-0.5) | -0.0227 | 0.0973 | 0.94 | -0.0631 | 0.0901 | 0.93 |
| $p_{acc}$ | 0.987 | | | | | |

**Table 2.** Simulation study results with misspecified models, including model(i): fitting a univariate model for each outcome, model (ii): assuming no latent class exist and implementing post-hoc clustering of fitted parameters, model (iii): assuming *K* = 2 latent classes and model (iv): assuming homogeneous convariance between classes.

| Model | WAIC | P<sub>acc</sub> |
| --- | --- | --- |
| True model | 80833.53 | 0.987 |
| Model (i) - Outcome 1 | 84419.02 | 0.954 |
| Model (ii) - Outcome 2 |  | 0.578 |
| Model (ii) | NA | 0.802 |
| Model (iii) | 83821.56 | 0.943 |
| Model (iv) | 85878.98 | 0.941 |

**Figure 1.**
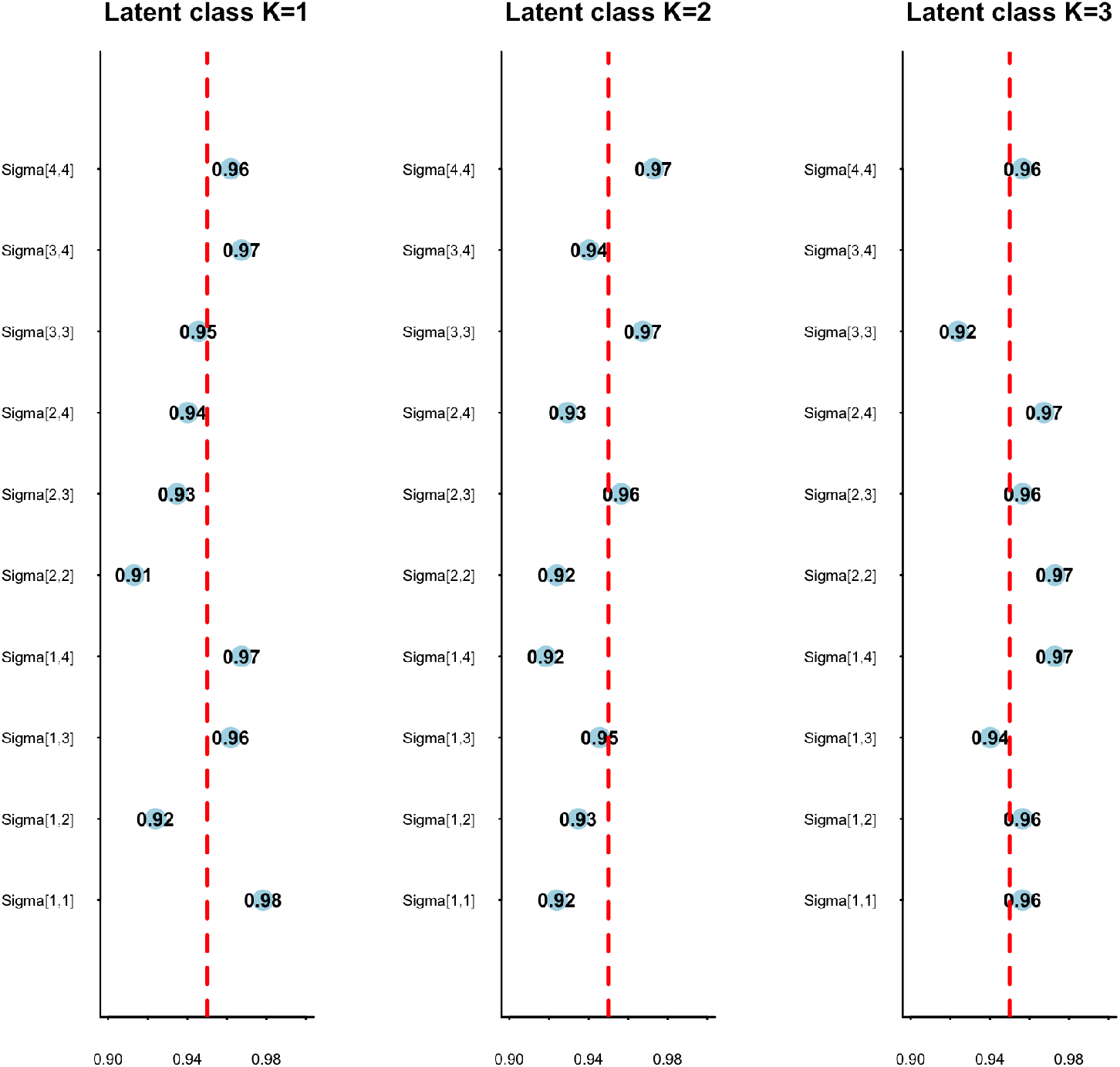
Coverage rate of the 95% credible intervals for estimated random effects covariance matrix for three latent classes.

### 4.2 Model misspecification performance

For the second part of simulation setting, we explored the model performance if it is misspecified. The true model was set simulated with *P* = 2 outcomes and *K* = 3 latent classes. The covariance matrix was assumed to be varied between classes. In addition to fitting the model with the true setting, we included the four misspecification cases, (i) Fitting a univariate model for each outcome (ii) assuming no latent class exist and implementing post-hoc clustering of fitted parameters, (iii) assuming *K* = 2 latent classes and (iv) assuming homogeneous convariance between classes. Classification accuracy *p*_*acc*_ will be computed for all three conditions of misspecification and compared. Additionally, WAICs were evaluated for the true model and misspecified cases (i), (iii) and (iv), which is an averaged estimates over 100 simulation samples. A combined WAIC of two univariate model was computed while each model provided an individual prediction accuracy value.

When the data are generated from *P* = 2 outcomes and *K* = 3 latent classes, the true model is most often preferred in terms of either WAIC (80833.53) or *p*_*acc*_ (0.987). The misspecification of the number of latent classes or covariance structure results in worse model performance with respect to the increasing WAICs. The univariate modeling shows an improved combined WAIC value compared with Model (iv) while presents a great discrepancy of prediction accuracy between two outcomes. The outcome 1, which were simulated with more separable random intercepts, greatly outperforms the outcome 2. Both model fitting performance and classification accuracy are improved with joint modeling with both outcomes, which supports our proposal of fitting multivariate model for ADNI data. Finally, the post-hoc classification based on fitted intercepts and slopes provides a poor *p*_*acc*_ compared with the latent class models, including those misspecified models.

## 5 Statistical Analysis

### 5.1 Data Structure

The BMGMM model was fit to the data from ADNI, mainly focused on Alzheimer’s Disease Assessment Scale Cognitive Subscale (ADAS-Cog) and MRI measures. The ADAS-Cog[25] is a cognitive assessment and higher scores indicate severer cognitive dysfunction, which is frequently used in pharmaceutical trials. In addition, MRI measures from multiple regions of interest (ROIs), including hippocampus, middle temporal lobe (Mid-Temp), fusiform, and entorhinal volumes[26, 27], are considered to be implicated in the pathophysiology of AD. As it is showed in Figure 2, the ADAS-Cog presents a more significant separation between three diagnostic stages, which is recorded at the entry of the study, while the MRI measures are harder to distinguish the stages. We fitted the BMGMM for these measures and predicted the latent stages of the AD progression.

**Figure 2.**
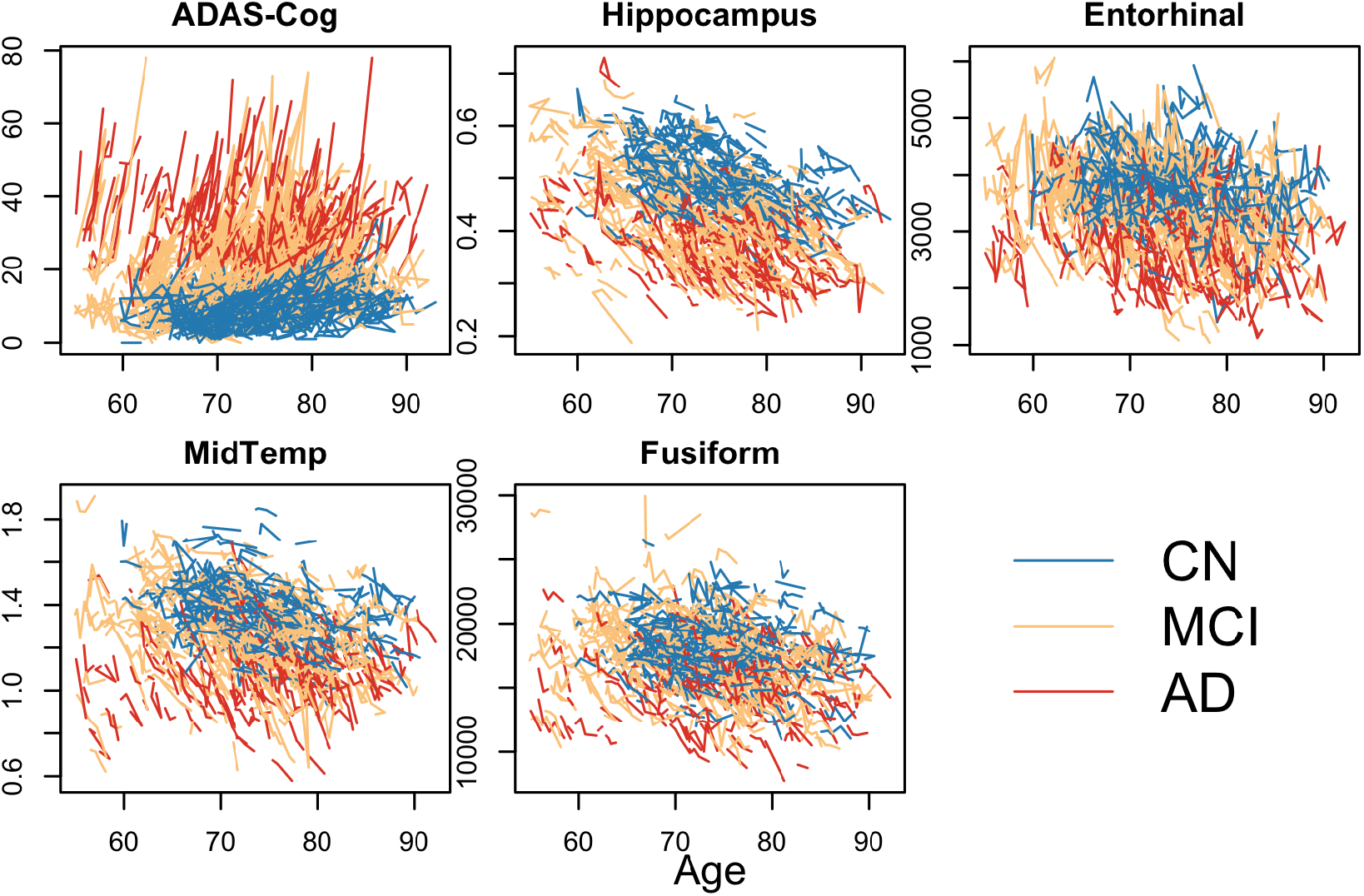
Spaghetti plots of the observed biomarker values of ADAS-Cog, hippocampus, middle temporal lobe, fusiform, and entorhinal volumes from 745 subjects in the Alzheimer’s Disease Neuroimaging Initiative with respect to their age over time. The colors indicate diagnostic stage at entry, including cognitively normal (blue), mild cognitive impaired (yellow) and dementia (red).

One goal of this analysis is to explore whether the predicted stages can inform both current and future development of the disease and provide sub-class classification with respect to the recorded diagnosis. With this consideration, the measurement Clinical Dementia Rating Sum of Boxes (CDRSB) was incorporated as a reference biomarker for indicating the progression of the measures in varying latent stages. In addition, we also evaluated the ability of our predicted latent states of indicating the change of diagnostic stages.

To achieve these goals, we fitted the BMGMM model including longitudinal measurements of ADAS-Cog and MRI measures of 745 subjects from the ADNI study. In accordance with three types of diagnostic stages at baseline (CN, MCI and dementia), we also assumed the each model with 3 latent classes (L1, L2 and L3). The outcome measures were standardized to be within range 0 and 1 for stability of fitting the covariance matrix. The carriage of the APOE*e*4 allele was used as the fixed effect covariate for each outcome and the covariates for class parameters included an intercept and the indicator of elevated amyloid. Meanwhile, based on the ground truth that the higher ADAS-Cog score indicates severer stages of disease, we assigned constrained order for the random intercepts of ADAS-Cog, i.e. larger random intercepts indicate higher latent classes. Two parallel Markov chains were run for 5000 iterations and the first 2500 warm-up iterations were discarded. The posterior mean and the 95% credible intervals were calculated using the obtained samples for each parameter.

### 5.2 Data analysis

Figure 3 shows the population- and subject-level predictions according to time after entry, colored with predicted latent states (left) and diagnostic stages in ADNI study (right). The predicted latent states present a clearer separation between each other compared with the diagnostic states, especially for estimation in MRI measures. The posterior means (95% credible intervals) for model parameters are given in Table 3. For ADAS-Cog and MidTemp volumes, all class-specific random effects of both intercepts and slopes illustrate a significant separation between three predicted latent states. As for the hippocampus and entorhinal volumes, the credible intervals of the estimated random intercepts and slopes between latent class 2 and 3 are overlapping, indicating that two latent class can be sufficient for modeling these two biomarkers. The fusiform volumes show a similar pattern in intercepts but different speed in slopes within latent class 2 and 3.

**Figure 3.**
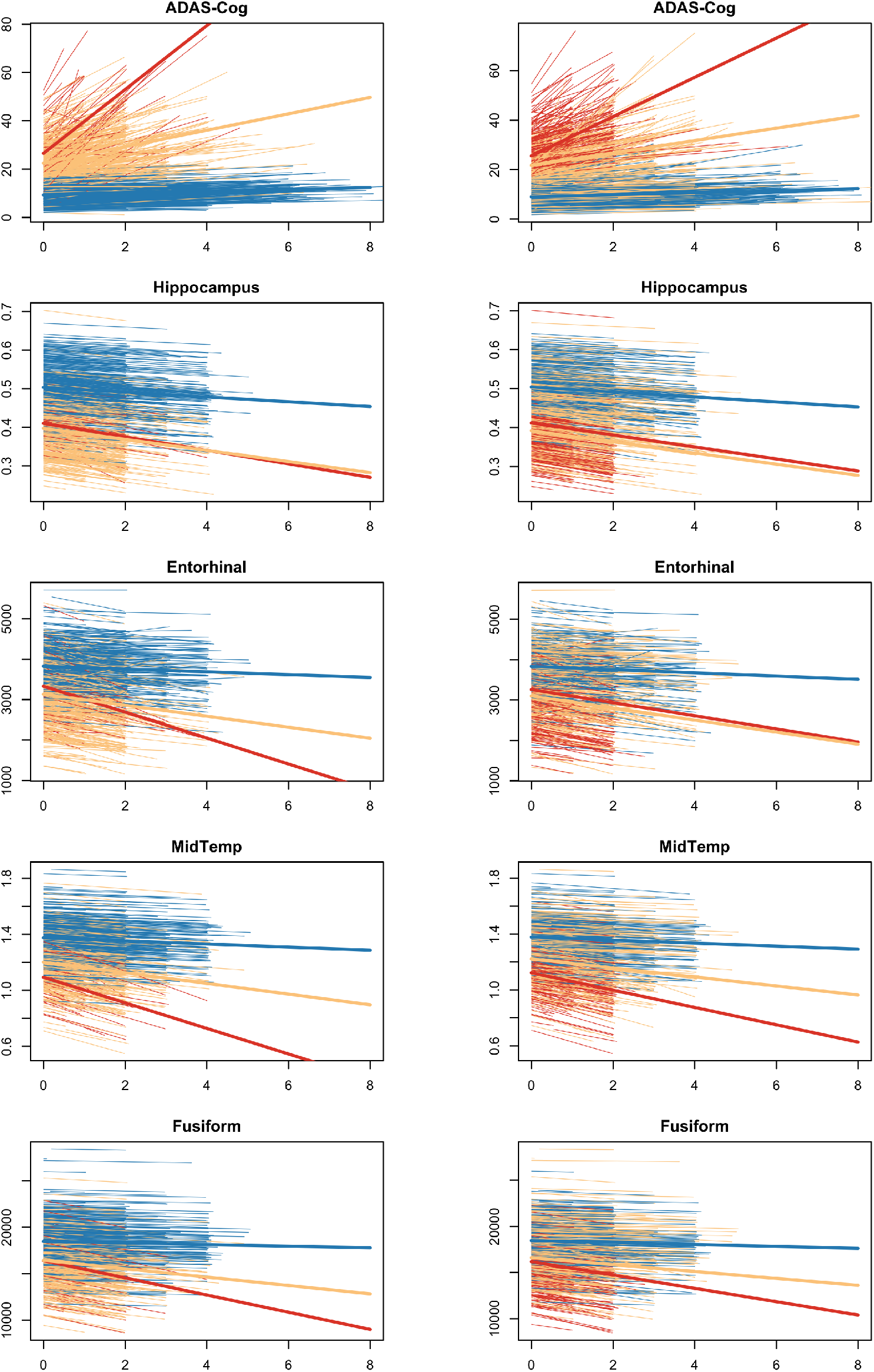
The modeled population and individual trajectories of ADAS-Cog, hippocampus, middle temporal lobe, fusiform, and entorhinal volumes (top to bottom). The colors indicate predicted latent classes (left panels) or diagnostic stages (right panels), including cognitively normal or latent class 1 (blue), mild cognitive impaired or latent class 2 (yellow) and dementia or latent class 3 (red).

**Table 3.**
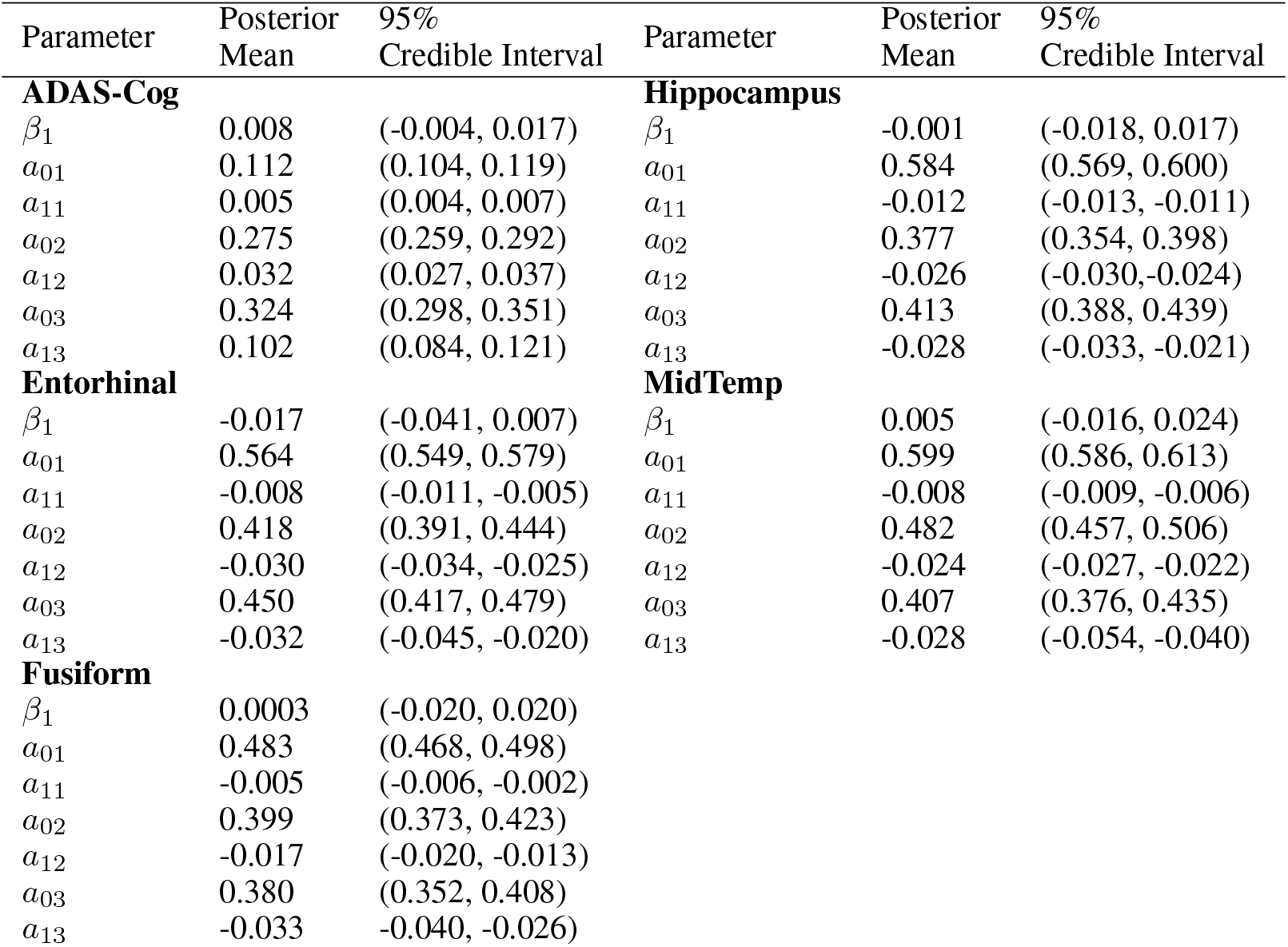
Results of posterior estimates of parameters for the proposed BMGMM fit to ADAS-Cog and four MRI measures.

| Parameter | Posterior Mean | 95% Credible Interval | Parameter | Posterior Mean | 95% Credible Interval |
| --- | --- | --- | --- | --- | --- |
| **ADAS-Cog** |  |  | **Hippocampus** |  |  |
| $\beta_1$ | 0.008 | (-0.004, 0.017) | $\beta_1$ | -0.001 | (-0.018, 0.017) |
| $a_{01}$ | 0.112 | (0.104, 0.119) | $a_{01}$ | 0.584 | (0.569, 0.600) |
| $a_{11}$ | 0.005 | (0.004, 0.007) | $a_{11}$ | -0.012 | (-0.013, -0.011) |
| $a_{02}$ | 0.275 | (0.259, 0.292) | $a_{02}$ | 0.377 | (0.354, 0.398) |
| $a_{12}$ | 0.032 | (0.027, 0.037) | $a_{12}$ | -0.026 | (-0.030, -0.024) |
| $a_{03}$ | 0.324 | (0.298, 0.351) | $a_{03}$ | 0.413 | (0.388, 0.439) |
| $a_{13}$ | 0.102 | (0.084, 0.121) | $a_{13}$ | -0.028 | (-0.033, -0.021) |
| **Entorhinal** |  |  | **MidTemp** |  |  |
| $\beta_1$ | -0.017 | (-0.041, 0.007) | $\beta_1$ | 0.005 | (-0.016, 0.024) |
| $a_{01}$ | 0.564 | (0.549, 0.579) | $a_{01}$ | 0.599 | (0.586, 0.613) |
| $a_{11}$ | -0.008 | (-0.011, -0.005) | $a_{11}$ | -0.008 | (-0.009, -0.006) |
| $a_{02}$ | 0.418 | (0.391, 0.444) | $a_{02}$ | 0.482 | (0.457, 0.506) |
| $a_{12}$ | -0.030 | (-0.034, -0.025) | $a_{12}$ | -0.024 | (-0.027, -0.022) |
| $a_{03}$ | 0.450 | (0.417, 0.479) | $a_{03}$ | 0.407 | (0.376, 0.435) |
| $a_{13}$ | -0.032 | (-0.045, -0.020) | $a_{13}$ | -0.028 | (-0.054, -0.040) |
| **Fusiform** |  |  |  |  |  |
| $\beta_1$ | 0.0003 | (-0.020, 0.020) | | | |
| $a_{01}$ | 0.483 | (0.468, 0.498) | | | |
| $a_{11}$ | -0.005 | (-0.006, -0.002) | | | |
| $a_{02}$ | 0.399 | (0.373, 0.423) | | | |
| $a_{12}$ | -0.017 | (-0.020, -0.013) | | | |
| $a_{03}$ | 0.380 | (0.352, 0.408) | | | |
| $a_{13}$ | -0.033 | (-0.040, -0.026) | | | |

Figure 4 displays the predicted individual-level progression intercepts and slopes from different model setting for ADAS-Cog and all four MRI measure. The raw slopes and intercepts were computed from linear regression of each standardized outcome trajectory individually and a post-hoc clustering was fitted on the estimated parameters via a multivariate Gaussian mixture model, assuming heterogeneous covariance structure across classes. These figures support our assumptions of the non-constant convariance among varying latent classes and concur with the results of class-specific parameters in Table 3, i.e. ADAS-cog and Midtemp volumes play a dominant role in separating the latent disease stages into three classes.

**Figure 4.**
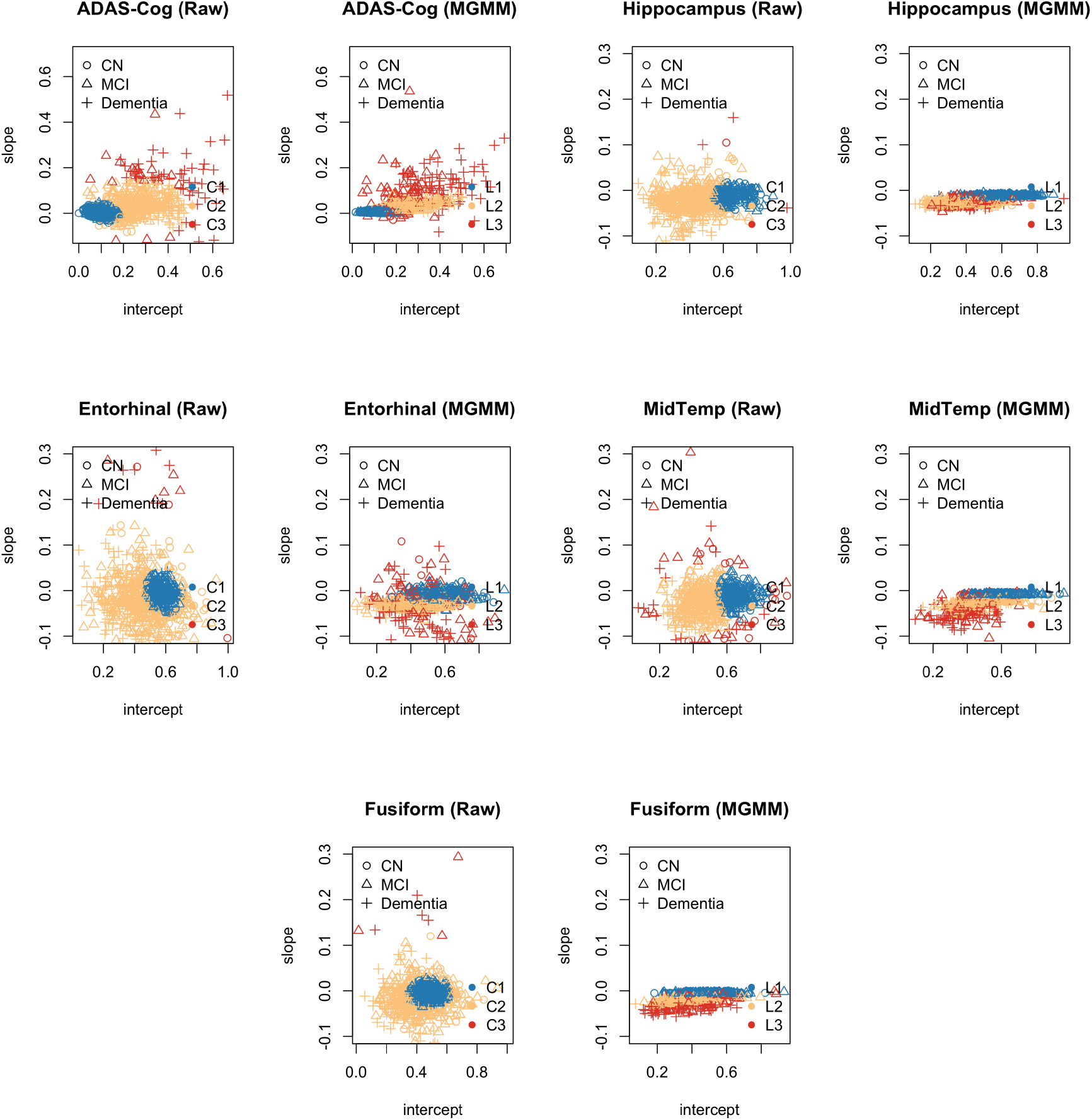
The distribution of estimated individual progression intercepts and slopes from raw estimation and multivariate GMM (right) for ADAS-Cog, hippocampus, middle temporal lobe, fusiform, and entorhinal volumes. The dots are colored with predicted latent states and symbolized with diagnostic states.

Table 4 provides the classification summary table between the diagnostic stages and predicted latent states. The posterior proportions indicate that almost all patients diagnosed with CN at baseline are classified with either L1 or L2 while for those MCI and dementia patients at baseline, most of them are classified as either L2 or L3. Combined with results showed in Figure 3, CN patients who are predicted in class L2 and MCI patients who are predicted in class L3 present a more severe pattern of disease progression, which illustrates potential sub-groups for these patients.

**Table 4.** Classification summary table for MGMM fitted with ADAS-Cog and four MRI measures.

| Diagnostic State | Predicted Latent Class |  |  |
| --- | --- | --- | --- |
|  | L1 | L2 | L3 |
| CN | 0.614 | 0.377 | 0.009 |
| MCI | 0.055 | 0.579 | 0.366 |
| Dementia | 0.106 | 0.465 | 0.429 |

Figure 5 further explores the varying patterns of disease development within different predicted latent states. The CDRSB measurements were incorporated as a reference biomarker and the slopes and intercepts were similarly calculated with linear regression of each standardized outcome trajectory individually. For MCI patients, almost all patients classified as L1 didn’t develop into dementia in follow-up diagnosis while for those who were classified as L2 or L3, a certain proportion of them appeared to be in worsen conditions. Furthermore, patients in L3 are most likely to deteriorate among all three latent states. Table 5 concludes the estimated model parameters of ADAS-Cog, entorhinal volumes and CDRSB within different disease stages and changing states, which conforms with our findings in Figure 5.

**Figure 5.**
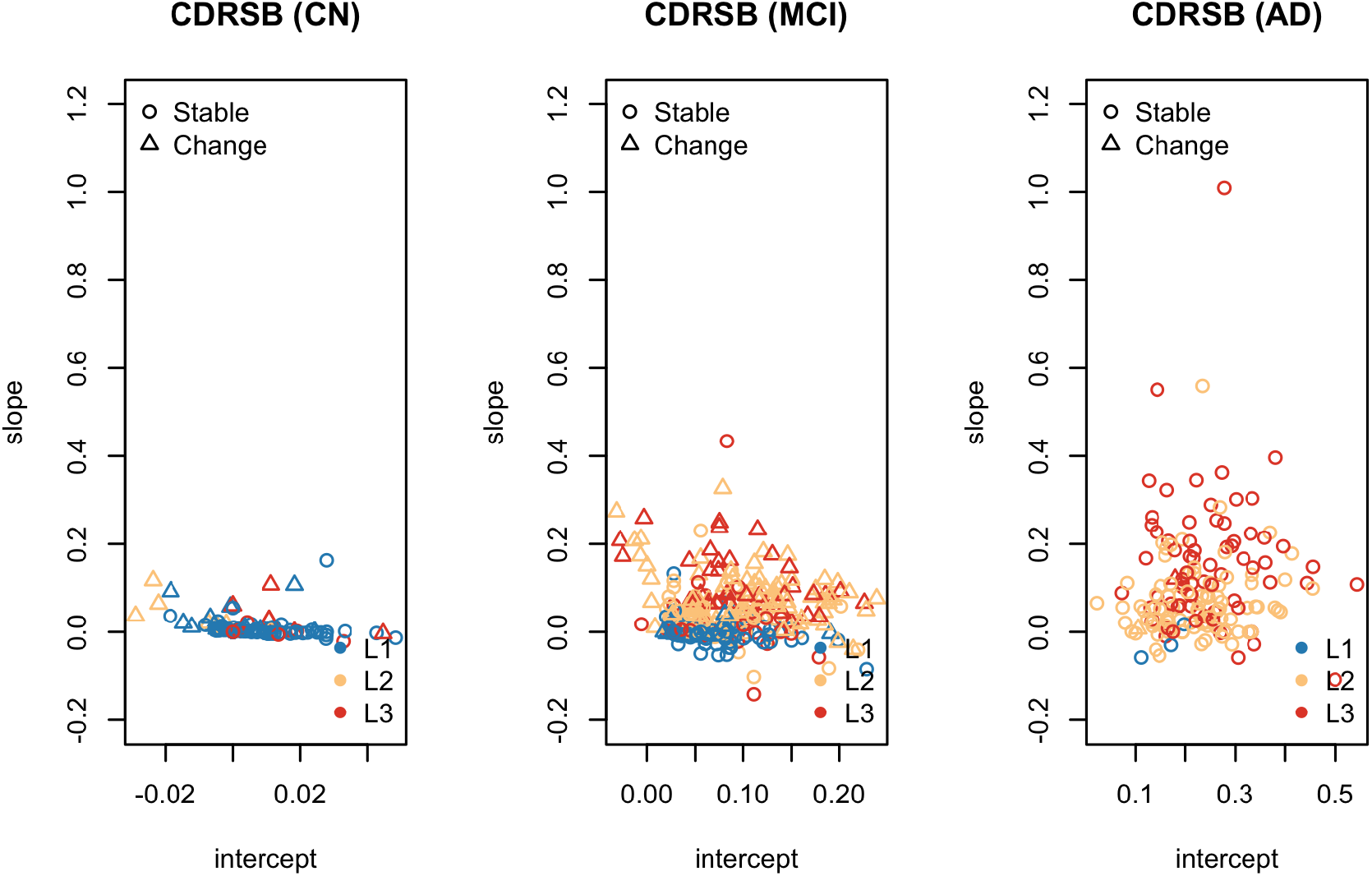
The individual-levle progression intercept and slope of CDRSB for CN(left), MCI(middel) and dementia(right), colored with predicted latent states. The symbol types represent whether the individual has developed to a severe stage during the study, i.e. CN - MCI (Dementia) or MCI - Dementia.

**Table 5.** Changing states summary table for ADAS-Cog, ventricular volumes and CDRSB.

| Parameter | ADAS-Cog |  |  | Entorhinal |  |  | CDRSB |  |  |
| --- | --- | --- | --- | --- | --- | --- | --- | --- | --- |
|  | CN | MCI | Dementia | CN | MCI | Dementia | CN | MCI | Dementia |
| Change = 1 |  |  |  |  |  |  |  |  |  |
| intercept | 0.151 | 0.256 | 0.337 | 0.541 | 0.451 | 0.418 | 0.001 | 0.099 | 0.143 |
| slope | 0.019 | 0.050 | 0.088 | -0.019 | -0.030 | -0.040 | 0.030 | 0.082 | 0.064 |
| Change = 0 |  |  |  |  |  |  |  |  |  |
| intercept | 0.104 | 0.171 | 0.366 | 0.559 | 0.525 | 0.382 | 0.003 | 0.071 | 0.234 |
| slope | 0.006 | 0.020 | 0.738 | -0.009 | -0.016 | -0.034 | 0.002 | 0.013 | 0.103 |

## 6 Discussion

We developed a Bayesian multiviariate growth mixture model for multivariate longitudinal data. The model is appealing for it incorporates several hierarchical approaches for generating parameters, including Half-t priors for standard deviations and Pólya–Gamma data augmentation for class-specific coefficients in multinomial regression. Compared with previous works with growth mixture models, these improvements not only promote the stability of model fitting involved with multiple outcomes and latent classes simultaneously, as well as function as a fully Bayesian hierarchical model, which is efficient and flexible.

The advantages of the proposed model have been supported with our simulation results. We showed that our model can be accommodated with varying combinations of latent classes and outcomes. For comparison purpose, we also implemented the simulation with commonly-used Inverse-Wishart priors for covariance matrix, which failed to converge in 50% simulation samples. In addition, the Bayesian auxiliary variable algorithm, introduced by Holmes and Held (2006)[19], was examined in binomial and multinomial sampling. Pólya–Gamma algorithm outperforms it both in efficiency and prediction accuracy, especially for multinomial cases. In addition, second part of the simulation experiments proved that the misspecification of the model can impair the model performance, validated with WAIC and prediction accuracy.

We applied our method to the ADNI data to assess the severity growth of ADAS-Cog and four MRI measures simultaneously and predict the latent disease states of individuals. We identified three clusters and compared them with the recorded diagnostic stages in the data. We found that CN participants are more likely to be included in latent class 1 and 2 while participants in mild dementia tend to be in latent class 2 and 3. In addition, MCI participants have more variability with the predicted states. Furthermore, we showed that our predicted latent states indicate the progression direction for individuals with different diagnostic stages, which is helpful in guidance with the AD development tracking in clinical studies.

Further works are required to extend current study. Firstly, more biomarker outcomes can be incorporated and compared for finding optimal combinations in predicting the latent disease stages of AD progression. Additionally, instead of using the linear model and normality assumption, other methods can be implemented to capture more variability within the data, such as splines and skew-normal distributions. Another potential direction is to include functional data analysis for modeling the random effects, which is flexible and able to picture deeper associations between the intercepts and slopes.

## References

[1] Clifford R Jack Jr, David S Knopman, William J Jagust, Leslie M Shaw, Paul S Aisen, Michael W Weiner, Ronald C Petersen, and John Q Trojanowski. Hypothetical model of dynamic biomarkers of the alzheimer’s pathological cascade. The Lancet Neurology, 9(1):119–128, 2010.

[2] Clifford R Jack Jr, David S Knopman, William J Jagust, Ronald C Petersen, Michael W Weiner, Paul S Aisen, Leslie M Shaw, Prashanthi Vemuri, Heather J Wiste, Stephen D Weigand, et al. Update on hypothetical model of alzheimer’s disease biomarkers. Lancet neurology, 12(2):207, 2013.

[3] Linda K McEvoy and James B Brewer. Biomarkers for the clinical evaluation of the cognitively impaired elderly: amyloid is not enough. Imaging in medicine, 4(3):343, 2012.

[4] Milton C Biagioni and James E Galvin. Using biomarkers to improve detection of alzheimer’s disease. Neurodegenerative disease management, 1(2):127–139, 2011.

[5] Ronald C Petersen. Mild cognitive impairment as a diagnostic entity. Journal of internal medicine, 256(3):183–194, 2004.

[6] Mei Sian Chong and Suresh Sahadevan. Preclinical alzheimer’s disease: diagnosis and prediction of progression. The Lancet Neurology, 4(9):576–579, 2005.

[7] Jorge L Bernal-Rusiel, Douglas N Greve, Martin Reuter, Bruce Fischl, Mert R Sabuncu, Alzheimer’s Disease Neuroimaging Initiative, et al. Statistical analysis of longitudinal neuroimage data with linear mixed effects models. Neuroimage, 66:249–260, 2013.

[8] Susan M Landau, Danielle Harvey, Cindee M Madison, Robert A Koeppe, Eric M Reiman, Norman L Foster, Michael W Weiner, William J Jagust, Alzheimer’s Disease Neuroimaging Initiative, et al. Associations between cognitive, functional, and fdg-pet measures of decline in ad and mci. Neurobiology of aging, 32(7):1207–1218, 2011.

[9] P Vemuri, HJ Wiste, SD Weigand, LM Shaw, JQ Trojanowski, MW Weiner, David S Knopman, Ronald Carl Petersen, CR Jack, et al. Mri and csf biomarkers in normal, mci, and ad subjects: predicting future clinical change. Neurology, 73(4):294–301, 2009.

[10] Michael C Donohue, Hélène Jacqmin-Gadda, Mélanie Le Goff, Ronald G Thomas, Rema Raman, Anthony C Gamst, Laurel A Beckett, Clifford R Jack Jr, Michael W Weiner, Jean-François Dartigues, et al. Estimating long-term multivariate progression from short-term data. Alzheimer’s & Dementia, 10:S400–S410, 2014.

[11] Bengt Muthén and Kerby Shedden. Finite mixture modeling with mixture outcomes using the em algorithm. Biometrics, 55(2):463–469, 1999.

[12] Robert H Pietrzak, Yen Ying Lim, David Ames, Karra Harrington, Carolina Restrepo, Ralph N Martins, Alan Rembach, Simon M Laws, Colin L Masters, Victor L Villemagne, et al. Trajectories of memory decline in preclinical alzheimer’s disease: results from the australian imaging, biomarkers and lifestyle flagship study of ageing. Neurobiology of aging, 36(3):1231–1238, 2015.

[13] Feng V Lin, Xixi Wang, Rachel Wu, George W Rebok, Benjamin P Chapman, Alzheimer’s Disease Neuroimaging Initiative, et al. Identification of successful cognitive aging in the alzheimer’s disease neuroimaging initiative study. Journal of Alzheimer’s Disease, 59(1):101–111, 2017.

[14] Jeannie-Marie S Leoutsakos, Sarah N Forrester, Christopher D Corcoran, Maria C Norton, Peter V Rabins, Martin I Steinberg, Joann T Tschanz, and Constantine G Lyketsos. Latent classes of course in alzheimer’s disease and predictors: the cache county dementia progression study. International journal of geriatric psychiatry, 30(8):824–832, 2015.

[15] Dongbing Lai, Huiping Xu, Daniel Koller, Tatiana Foroud, and Sujuan Gao. A multivariate finite mixture latent trajectory model with application to dementia studies. Journal of applied statistics, 43(14):2503–2523, 2016.

[16] Alan Huang, Matthew P Wand, et al. Simple marginally noninformative prior distributions for covariance matrices. Bayesian Analysis, 8(2):439–452, 2013.

[17] Nicholas G Polson, James G Scott, and Jesse Windle. Bayesian inference for logistic models using pólya–gamma latent variables. Journal of the American statistical Association, 108(504):1339–1349, 2013.

[18] Ronald Carl Petersen, PS Aisen, Laurel A Beckett, MC Donohue, AC Gamst, Danielle J Harvey, CR Jack, WJ Jagust, LM Shaw, AW Toga, et al. Alzheimer’s disease neuroimaging initiative (adni): clinical characterization. Neurology, 74(3):201–209, 2010.

[19] Chris C Holmes, Leonhard Held, et al. Bayesian auxiliary variable models for binary and multinomial regression. Bayesian analysis, 1(1):145–168, 2006.

[20] Sylvia Frühwirth-Schnatter and Helga Wagner. Stochastic model specification search for gaussian and partial non-gaussian state space models. Journal of Econometrics, 154(1):85–100, 2010.

[21] Andrew Gelman et al. Prior distributions for variance parameters in hierarchical models (comment on article by browne and draper). Bayesian analysis, 1(3):515–534, 2006.

[22] Sumio Watanabe. Asymptotic equivalence of bayes cross validation and widely applicable information criterion in singular learning theory. Journal of Machine Learning Research, 11(Dec):3571–3594, 2010.

[23] Matthew Stephens. Dealing with label switching in mixture models. Journal of the Royal Statistical Society: Series B (Statistical Methodology), 62(4):795–809, 2000.

[24] Panagiotis Papastamoulis. label. switching: An r package for dealing with the label switching problem in mcmc outputs. arXiv preprint arXiv:1503.02271, 2015.

[25] Wilma G Rosen, Richard C Mohs, and Kenneth L Davis. A new rating scale for alzheimer’s disease. The American journal of psychiatry, 1984.

[26] Iryna Lobanova and Adnan I Qureshi. The association between cardiovascular risk factors and progressive hippocampus volume loss in persons with alzheimer’s disease. Journal of vascular and interventional neurology, 7(5):52, 2014.

[27] Clifford R Jack Jr, Heather J Wiste, Stephen D Weigand, Terry M Therneau, Val J Lowe, David S Knopman, Jeffrey L Gunter, Matthew L Senjem, David T Jones, Kejal Kantarci, et al. Defining imaging biomarker cut points for brain aging and alzheimer’s disease. Alzheimer’s & Dementia, 13(3):205–216, 2017.

